# Spatially-explicit genomics of *An. gambiae s.l* uncovers fine-scale population structure and mechanisms of insecticide resistance

**DOI:** 10.1101/2025.06.13.655782

**Authors:** Sanjay C Nagi, John Essandoh, Nicholas Ato-Egyir, Eric R Lucas, Alistair Miles, Kwame Desewu, Ignatius Williams, Alexander Egyir-Yawson, Luigi Sedda, David Weetman, Martin J Donnelly

## Abstract

Progress in malaria control in sub-Saharan Africa is stalling, partly due to the spread of insecticide resistance in *Anopheles* vectors. Monitoring the evolution of insecticide resistance alleles and their spatial heterogeneity is important for malaria control programmes, and genomic surveillance has emerged as a pivotal tool for this purpose. Earlier genomics research has typically employed convenience-based sampling, and research has yet to be performed to optimise sampling regimens for malaria vector genomics. In an earlier study, we developed a spatially explicit sampling framework that considers the underlying ecology to enable sampling mosquitoes with reduced bias. We applied this framework to sample and perform whole-genome sequencing on 485 individual specimens of *An. gambiae s.l* mosquitoes from Obuasi, central Ghana, an area with extensive gold mining activities. In this region, *An. gambiae s.l* have been documented as highly resistant to multiple insecticides, including pyrethroids used for the treatment of bed-nets. High resolution of fine-scale population structure enabled the detection of isolation-by-distance in *An. coluzzii* in Obuasi, whilst finding that at this scale, geographic distance, rather than the underlying habitat, drives population structure. We examine methods to estimate mosquito kinship, demonstrating that polymorphic chromosomal inversions significantly confound established analytical tools, while revealing how the mosquito’s chromosome architecture generates high variance in genetic relatedness among kin. Using genome-wide selection scans and clustering, we discover the novel mutations *Gste2-*F120L and *Cyp9K1*-N224I in detoxification enzymes that are driving selective sweeps, likely in response to insecticidal pressure. We find that the frequencies of some resistance variants are associated with artisanal-gold mines, and elucidate the continued evolution of the *Voltage-gated sodium channel*, the target of pyrethroid insecticides. Overall, we show that sampling vectors strategically may enhance our ability to perform effective genomic surveillance.

## Introduction

Evidence suggests insecticide resistance may be compromising the efficacy of malaria control interventions^1,2^. Repeated exposure to toxic chemicals has led to rapid adaptation in mosquito vectors, allowing them to survive even extreme doses of insecticide^3^. Recent developments in genomic technologies have facilitated the study of the genetic basis of insecticide resistance^4–10^. Although certain resistance alleles, such as the well-known knockdown resistance (*kdr)* mutations, have spread or arisen independently in *Anopheles* across much of sub-Saharan Africa^7^, many other mechanisms of resistance seem to be localised to specific areas^9,11^. This may result from heterogeneities in insecticide usage resulting in hotspots of selection pressure^12^. It could also arise from micro-spatial population structure, with minor barriers to gene flow slowing the spread of new mutations, particularly if their selective advantage is relatively small. Understanding these processes will be important for sustainable, targeted vector control, where tailored interventions can be prioritised in hotspots of malaria transmission^13^.

The majority of genomic studies have taken place over extremely large spatial scales, often continent-wide, leaving researchers unable to explore questions on micro-spatial scales^6^, and providing limited information to inform control. They have also been performed without predefined sampling frameworks - sampling sites have typically been selected for their convenience and familiarity, rather than being designed to maximise informational value. Sedda *et al*. developed a sampling framework for the surveillance of malaria mosquitoes, which incorporates ecological data to optimise the accuracy of mosquito distribution estimates^14^. In brief, the framework combines a regular lattice with random points as close pairs, to maximise spatial coverage, the representativeness of ecological zones and vector spatial autocorrelation. In this framework, 70% of sampling points are in a grid, with the remaining 30% of points randomly allocated as close-pairs. To ensure ecological representativeness, each ecological stratum contains a number of sampling sites proportional to the stratum size. The framework, by ensuring representative sampling across habitats, can improve understanding of how genetic variation is distributed among ecological niches and how environmental variation influences the evolution and spread of insecticide resistance. Understanding conduits and barriers to gene flow is also central to the use of transgenic gene drive technologies, designed for either population replacement or suppression^15,16^.

In this study, we collected *An. gambiae s.l* mosquitoes using this ecologically-informed sampling framework in a 2,463 km^2^ region around Obuasi, central Ghana. Obuasi is located in the southern Ashanti region, known for substantial gold mining activity^17^. Mining activities are a mixture of large mines run by multinational corporations or artisanal and small-scale mines (ASM). ASMs have been reported to create *Anopheles* breeding sites^18^. Across Ghana and in Obuasi specifically, *An. gambiae s.l* have been documented to be highly resistant to a range of insecticides^9,19–21^. We perform a population genomic analysis using whole-genome sequencing of 485 *An. gambiae* mosquitoes, investigating relatedness and population structure at ultra-fine spatial scales. We explore the genomics of insecticide resistance and report allele frequencies of known insecticide resistance loci, detecting signals of selection and mutations driving selective sweeps.

## Results

### Sample collections and whole-genome sequencing

We collected mosquitoes indoors using CDC light traps from four houses at each sampling village in the Obuasi area, in a 9 week period from October to December 2018. A total of 1,859 *An. gambiae* s.l. mosquitoes were collected from 99 of 120 houses across 30 villages. Sampling was performed in the remaining 21 houses, but no mosquitoes were captured. Across all collections, the maximum number of *An. gambiae s.l* collected in a single night was 64, whilst the average nightly catch per house was 3.1. Supplementary Figure 1 shows the number of collected *An. gambiae s.l* per house over the sampling period. To select a subset of 500 specimens for sequencing we chose samples from all positive houses as evenly as possible, taking every mosquito from each house up to a maximum of 17 per house.

A total of 485 *An. coluzzii* (n=422) and *An. gambiae s.s.* (n=63) samples passed pre-sequencing QC and were whole-genome sequenced to an estimated coverage of 30X (Supplementary Table 1). No *An. arabiensis,* hybrids, or cryptic species were detected in the data. Reads were aligned to the AgamP4 reference genome and analysed as previously described^6^. We discovered a total of 43,730,731 segregating SNPs that passed our quality filters^6^. Figure 1 shows a map of the Obuasi region, including ecological stratification, sample collection sites, the relative proportion of *An. gambiae s.s* and *An. coluzzii* shown, and whether ASMs are present at the site.

**Figure 1.**
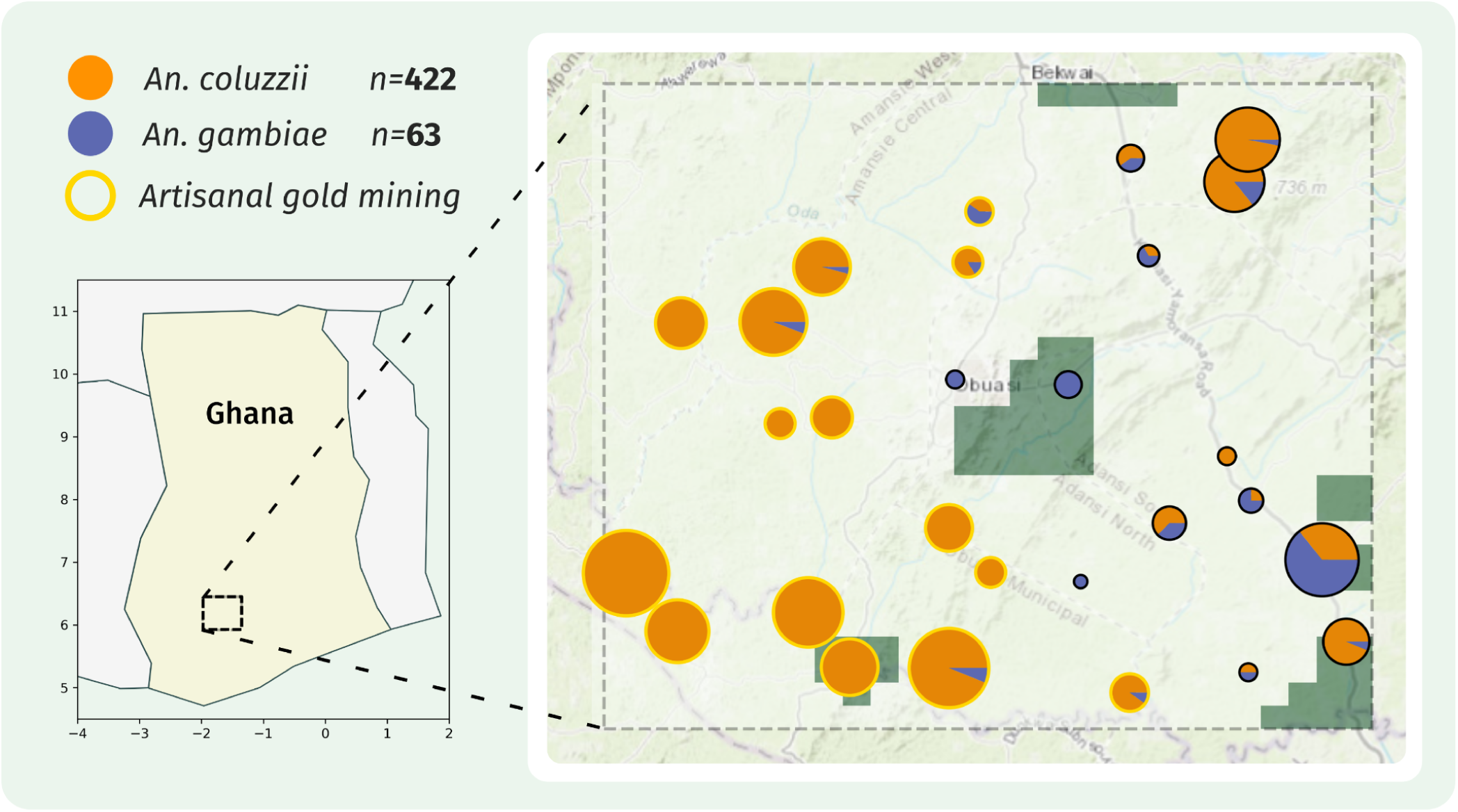
Sample map. A map of sampling locations from the Obuasi region, overlaid onto the ecological classifications from Sedda et al., 2019. The number of *An. coluzzii* and *An. gambiae* that passed sequencing quality control are shown, and the species composition at each site displayed as a pie chart. Sample collection sites which are the location of Artisanal gold mining occurs are highlighted with a gold circle. Green polygons = forest; rest of the area is a mixture of urban, wetland and grassland with similarities in evapotranspiration, elevation, and surface temperature.

### Population structure

To explore overall population structure in the data, we built a neighbour-joining tree from biallelic SNPs from a region of the 3L chromosomal arm between 15 Mb and 44 Mb, selected to avoid chromosomal inversions, selective sweeps and heterochromatin (Figure 2A), which may obscure underlying patterns of genetic differentiation between cohorts. The neighbour-joining tree showed genetic separation between *An. coluzzii* and *An. gambiae*, but there was no evidence for within-species population structure.

**Figure 2.**
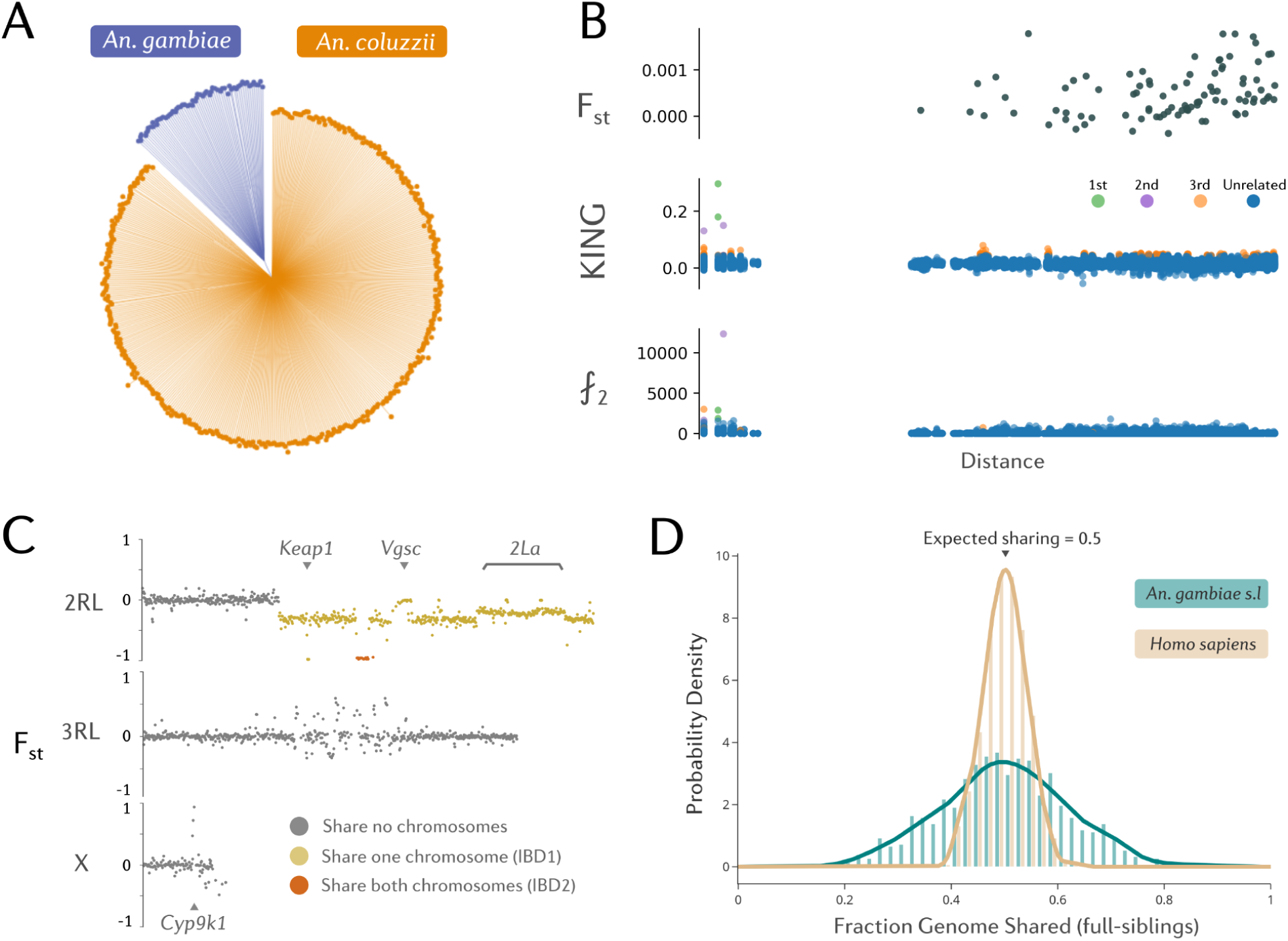
Population structure and isolation-by-distance. A) An unrooted neighbour-joining tree computed from pairwise genetic distances using biallelic SNP variants from the 3L chromosomal arm (3L:15,000,000-44,000,000). B) Measures of genetic distance vs the natural logarithm of geographic distance in kilometres. F_ST_ is calculated between sample collections sites for *An. coluzzii*. KING is a measure of genetic relatedness based on linkage disequilibrium. ⨏_2_ variants are mutations only found in two individuals and are a proxy for rare variants. The y-axis indicates the number of doubletons shared between pairs C) Genome-wide Hudson F_ST_ calculated between a pair of *An. coluzzii* individuals identified as third-degree kin, found 6.99 Km apart, in Domenase and Nkotumso. D) A probability density plot of the fraction genome shared in full-siblings, from simulations with ped-sim.

We performed principal components analysis (PCA) on the same genetic region (Supplementary Figure 2). As expected there is clear segregation by species, with PC1 capturing variation that distinguishes the two species, we also observe two outliers assigned as *An. gambiae s.s*. We also perform PCA for both species in isolation (Supplementary Figure 2). In the *An. gambiae* PCA, we observe the same two samples as outliers again (Supplementary Figure 2C). The removal of these two samples resulted in a PCA showing a lack of clear structure (Supplementary Figure 2D). Overall, the neighbour-joining tree and principal components analyses suggest that there is little differentiation into distinct subpopulations within *An. coluzzii* or *An. gambiae s.s*, in agreement with earlier findings that this region of West Africa is relatively homogeneous^6^.

The presence of pairs of outliers in the PCA analyses suggested that close kin may exist in our data. To estimate relatedness between samples, we calculated the KING-robust statistic^22^ from biallelic SNP markers across the whole-genome for every pair of individual mosquitoes in the dataset, using the software NgsRelate^23^. Initially, this resulted in within-species KING values which were tri-modal, which we determined to result from the presence of large, polymorphic chromosomal inversions in our data (Supplementary Text 2). Chromosomal inversions in *Anopheles gambiae* are known to be ancient - for example, the 2La inversion predates speciation into the *gambiae* complex, and therefore genetic differentiation between opposing karyotypes is very large - greater than the level of differentiation found between sibling species^4^. We therefore excluded the 2La and 2Rb inversion regions from our relatedness analyses.

Two full-sibling pairs were detected in the data. Both pairs were *An. coluzzii,* designated as full siblings (the expectation of KING value for full siblings is 0.25), and were sampled from Odumto in houses approximately 111m away from each other. The inferred kin pair found furthest apart were collected in Nkotumso and Domenase, 6.99 km away from each other, and predicted to be third-degree kin. To investigate genome-wide inheritance in detail and confirm kinship, we explored genome-wide patterns of genetic differentiation between the putative third-degree relatives. Based on genome-wide F_st_ (Figure 2C), this pair of *An. coluzzii* appear related, sharing a large IBD segment on chromosome 2. Three levels of F_st_ can be observed, depending on whether pairs share zero (F_st_ ≈ 0), one (IBD1, F_st_ ≈ -0.25), or both (IBD2, F_st_ ≈ -1) chromosomal segments from their ancestors at any given position. Supplementary Figure 5 shows F_st_ between the two individuals with the highest KING value, F_st_ on the X chromosome is -1, indicating that both individuals received the only paternal X chromosome, and the same maternal X chromosome.

Using the previously published^24^ estimates of the per-base recombination rate in *An. gambiae* of 10^-8^bp^-1^, we calculated the expected number of recombination events per generation for chromosome 2 using a binomial distribution. The expected value was 1.11 recombination events (95% CIs: 0-4) with 33% of generations having 0 recombination events (although this calculation does not account for crossover interference). This is fewer than the 2-3 events per chromosome observed in humans^25^, although most human chromosomes are much larger than *An. gambiae* chromosomes. *An. gambiae* and other mosquitoes also only have 3 chromosomes compared to humans (n=23 pairs). Thus, while the expected genetic relatedness between siblings of *An. gambiae* is 0.5, the variance around this value will be higher than for human siblings, and in some cases pairs might range from appearing more like genetic cousins, or closer to genetic twins. To explore this further, we simulated 10 replicate pedigrees of *An. gambiae* and *Homo sapiens* with ped-sim v1.4^26^, generating 800 individuals in total including 920 full-sibling pairs per species. Ped-sim uses a genetic map containing genome-wide recombination rates to simulate the propagation and inheritance of IBD segments in the pedigree. We then used these IBD segments to calculate the fraction of the genome shared between full-siblings (Figure 2D), demonstrating that in comparison to humans, *An. gambiae* kin have a much higher variance in genetic relatedness. This is likely to make kinship inference challenging in *An. gambiae*, and may mean that we should use an estimate of kinship likelihood rather than a single prediction of kinship category, particularly in future statistical methods relating to close-kin mark-recapture studies.

### Isolation by distance

Isolation-by-distance (IBD) is an important parameter in population genetics, describing the tendency for organisms closer in space to be more similar to each other, arising from distance-limited dispersal^27^. In West Africa, studies over a much broader geographical scale have found different rates of IBD when comparing *An. gambiae s.s* or *An. coluzzii^6^*, with *An. coluzzii* displaying stronger IBD, suggesting lower dispersal. Although we could not compare between species due to limited sample sizes in *An. gambiae s.s,* we performed an analysis of isolation-by-distance in *An. coluzzii*, at a much finer scale than has been previously attempted.

We calculated Hudson’s F_st_ between individuals from each sampling site. F_st_ was generally low, as was expected from the proximity of sampling locations. Using the pairwise F_st_ estimates between sample sites, we plotted linearised F_st_ against the geographic distance between sites (Figure 2B), and fitted a regression line as in the method of Rousset^28^. We initially excluded sampling locations with fewer than 10 mosquitoes, which resulted in 13 sampling locations in total. We observe a positive and significant slope of the regression line (p=0.0063), suggesting that genetic differentiation does increase with geographic distance in our dataset. Whilst isolation-by-distance has been observed between *An. gambiae s.l* populations before, the detection of isolation by distance within this dataset is novel given the micro-spatial sampling design. As another measure of population structure, we identified doubleton (⨏_2_) variants, alleles which are only found twice in the dataset and are thought to be indicative of recent mutational events as they have not yet had chance to drift to appreciable frequencies^29^. It was shown by Matheison and McVean that these loci provide a powerful means of detecting fine-scale population structure. As with F_st_, we found a significant association with the number of shared doubletons between two individuals and with geographic distance (p=4.5e-120). We also find that the individuals with the highest number of shared doubletons also tend to be siblings (Supplementary Figure 6).

To investigate any effect of environment on isolation-by-distance, we also gathered 13 ecological variables which were used to inform the sampling design^14^) (e.g. vegetation and land cover, elevation, temperature and precipitation). We removed correlated ecological variables, and calculated an overall ecological distance between each sample site, using Euclidean distance. We then performed partial Mantel tests on the physical distance, ecological distance and F_st_ matrices, to test for associations between two variables whilst controlling for a third^30^. When we tested for associations between geographic distance and F_st_ whilst controlling for ecology, the result was significant (p=0.0005). However, the reverse was not true - ecological distance did not have a significant effect on genetic distance when controlling for geographic distance (p=0.4893), suggesting that geographic distance drives differentiation at this scale, rather than isolation by environment. We similarly tested for potential genetic structuring due to the presence of artisanal gold mines; geographic distance was significantly associated with genetic distance when controlling for the presence of ASMs (p=0.006), but ASMs were not associated with genetic distance when controlling for geographic distance (p=0.6899), suggesting that ASMs are not a driver of population structure in Obuasi.

Supplementary table 2 shows karyotype frequencies for each sample site in Obuasi for the 2La and 2Rb inversions, which have previously been linked to ecological variation^31^. The 2La inversion was at 40.1% frequency in the dataset as a whole, at 41.1% in ecotype one, and 34.2% in ecotype two. We tested for differences in the frequency of 2La between the two ecotypes, however, this was not significant (Ꭓ^2^, p=0.31).

### The genomics of insecticide resistance

Given the high intensities of insecticide resistance in Ghana^19^, we investigated known insecticide resistance alleles and examined signatures of selection across the genome. We identified regions of the genome that are under selection with the H12 statistic (Figure 3), a measure of haplotype homozygosity capable of detecting both soft and hard selective sweeps^32^. To determine if any of these selective sweeps are shared between *An. gambiae* and *An. coluzzii*, we applied the H1X statistic, which measures the probability that any two haplotypes sampled from each cohort are identical in a given genomic window^33^. We also calculated frequencies of amino-acid mutations, amplifications or deletions (copy number variants) around insecticide resistance-associated loci (Supplementary Figure 7). Specific copy number variant alleles are reported in Supplementary Table 3. To further explore the haplotype structure at resistance loci whilst incorporating amino acid and copy number variation, we performed hierarchical clustering on diplotypes^34^. Diplotypes are stretches of diploid genotypes, sometimes referred to as multi-locus genotypes. We perform clustering on diplotypes instead of haplotypes as it allows us to resolve multiallelic SNPs and copy number variants, both of which are challenging to phase onto haplotypes.

**Figure 3.**
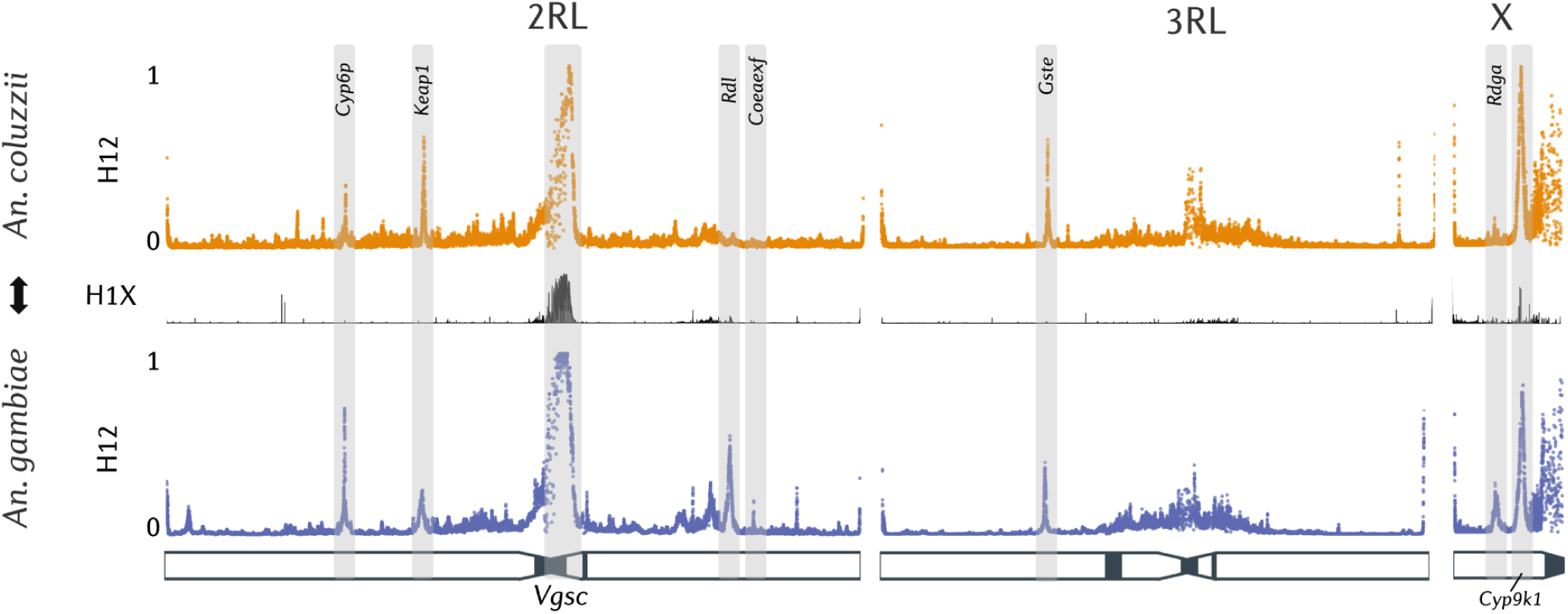
Genomics of insecticide resistance. H12 Genome-wide selection scans in *An. gambiae* and *An. coluzzii* from Obuasi detecting regions of the genome under positive selection. These scans are separated by a H1X introgression scan, a statistic which measures the probability of a haplotype being shared between two cohorts. A schematic of the *An. gambiae* PEST chromosomes is shown beneath the scans, with heterochromatin highlighted in black.

Supplementary Figure 6 shows allele frequencies of SNP variants in the *Vgsc, Ace1 and Gste2* genes and CNV variants at metabolic detoxification loci split by species. Overall, only the single *An. gambiae* population from New Edubiase is clearly distinct, with sampling villages showing little overall variation in allele frequencies (AMOVA, P value > 0.9).

In *An. coluzzii,* the *Vgsc-*995F mutation remains at high frequencies ranging between 76% and 94%. The remaining non-995F alleles at each site correspond to the V402L/I1527T haplotype. Recent data has shown that this V402L/I1527T haplotype is increasing in frequency and geographic range^7,35^, with high frequencies of the haplotype as far east as Chad, Niger (>50% AF) and even Kenya (∼38%)^36,37^, suggesting that the haplotype first discovered in Guinea, Burkina Faso, and Ghana, has already spread across the range of *An. coluzzii. Anopheles gambiae s.s* remains fixed for 995F haplotypes^7^. Between *An. gambiae s.s* and *An. coluzzii*, we can also observe distinct patterns of secondary *Vgsc* mutations occurring alongside L995F. In *An. coluzzii*, R254K, T791M, P1874S and I1940T are all found at low to moderate frequencies. These alleles are known to have arisen on the background of 995F haplotypes and might increase the resistance level or reduce fitness costs associated with resistance. In *An. gambiae s.s*, the secondary mutations are T791M, N1570Y, A1746S, I1853I, I1868T, and P1874L, in agreement with recent studies from West Africa^9,38^. Diplotype clustering confirms these patterns, demonstrating three broad diplotype clusters - a cluster homozygous for V402L and I1527T, a cluster homozygous for L995F, and a cluster heterozygous for both mutations (Supplementary Figure 8). The large H1X signal at this locus indicates that there are shared haplotypes between the two species, and indeed, the L995F cluster contains individuals of both *An. gambiae* and *An. coluzzii*, confirming earlier findings of adaptive introgression at this locus^39^.

There is a large contrast in the frequency of the *Ace1-*G280S mutation between species - 37% in *An. gambiae* compared to 1% in *An. coluzzii.* In *An. gambiae*, we observe a much higher prevalence of *Ace1* amplifications (62%) than the *Ace1-280S* mutation itself (37%). It is known that CNVs in the wild pairs wild-type Glycine alleles with resistant Serine alleles^8^. Samples that are homozygous for these CNVs can appear heterozygous for *Ace1*-280S, despite it being present on both chromosomes. Diplotype clustering again confirms the presence of haplotype sharing between *An. gambiae* and *An. coluzzii* at this locus, with a small number of *An. coluzzii* individuals residing within the *An. gambiae* G280S containing cluster (Supplementary Figure 9).

At the *Cyp6aa/p* gene cluster, a known metabolic resistance locus, we find the recently reported, resistance-predictive, *Cyp6p3-*E205D mutation in *An. gambiae^40^*, accompanied by little to no involvement of CNV alleles (Supplementary Figure 10). In *An. coluzzii,* however, we can observe multiple CNV alleles at appreciable frequency, including an allele which covers *Cyp6p15p*-*Cyp6p1*, another which covers *Cyp6aa1* and *Cyp6aa2*, and another from *Cyp6aa1*-*Cyp6p15p*. Each of these three CNV alleles seems to be driving its own selective sweep. The *Cyp6p3-*D155N allele is found on a swept diplotype which harbors no CNVs, and may be a promising new candidate for a causative role in resistance. At *Cyp9K1*, we find no evidence of haplotype sharing between *An. gambiae* and *An. coluzzii* (Figure 4A). Despite most *An. coluzzii* diplotypes belonging to a single swept cluster, there is no evidence for any amino acid variation in this species nor copy number variants. This suggests that a different mechanism is driving this sweep, such as a nearby genomic insertion. In agreement with the observations from diplotype clustering, we see a 79% frequency of *Cyp9k1* gene amplifications in *An. gambiae s.s* but low frequencies of *Cyp6aa/p* CNVs (<6%), with the opposite pattern observed in *An. coluzzii* (supplementary Figure 7). We can observe an amino acid variant, *Cyp9k1-*N224I, which looks like it is driving a selective sweep in *An. gambiae* (Figure 4A).

**Figure 4.**
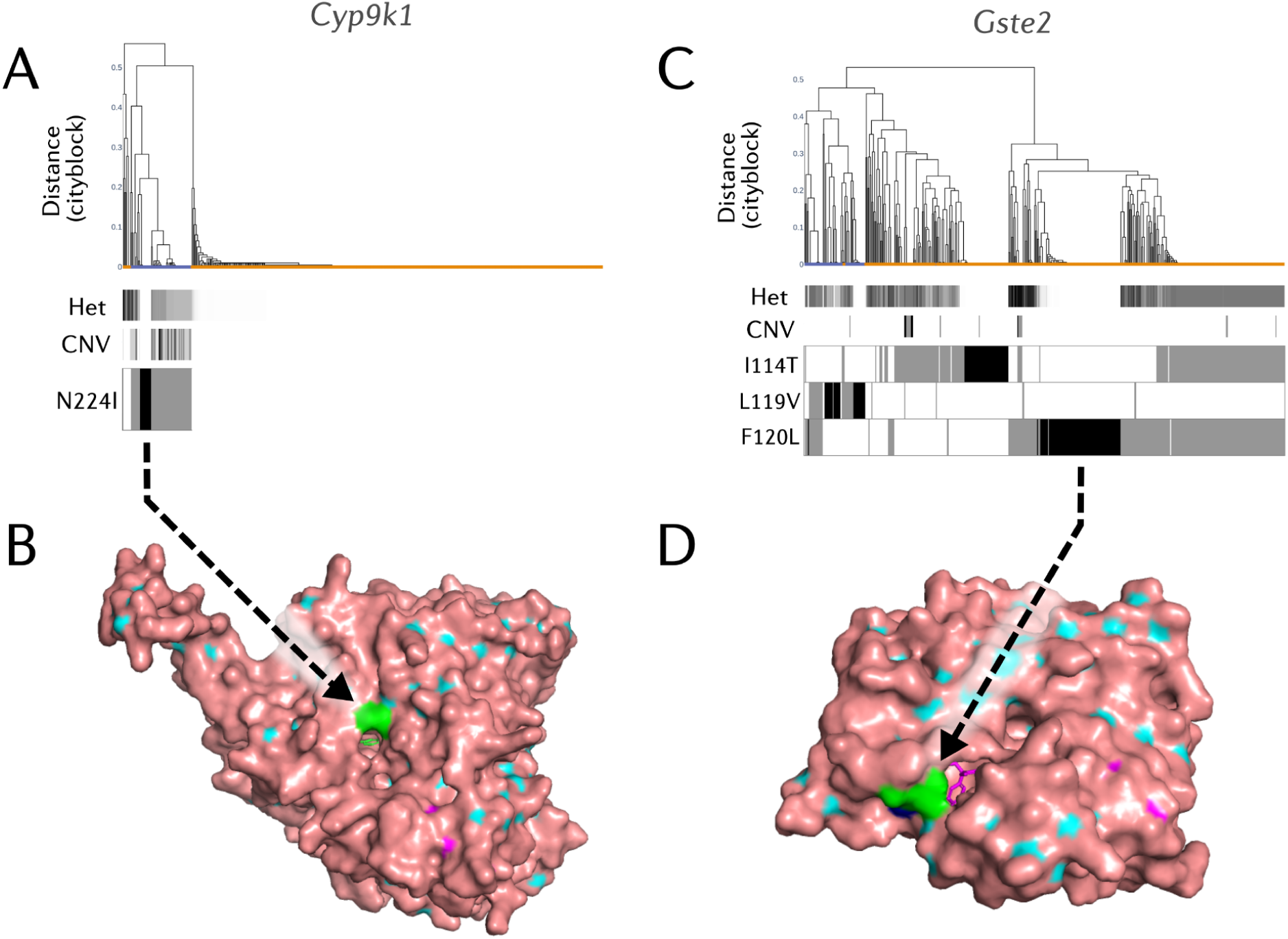
Resistance variant analysis. A) Diplotype clustering at Cyp9k1. We calculate pairwise distances between diplotypes. Each column in the figures is a diplotype ordered in the dendrogram by hierarchical clustering, using genetic distance based on city-block (Manhattan) distance and complete linkage. The leaves of the dendrogram are coloured by the species of the individual to which they belong. Note that due to overlapping points, not all dendrogram leaves can be seen. Underneath the dendrogram, the heterozygosity and CNV copy number of an individual are displayed as horizontal bars. Heterozygosity; individual-level heterozygosity was calculated as an average over all SNPs in the locus. Clusters with low sample heterozygosity or inter-sample genetic distances of zero are indicative of a selective sweep. CNV copy numbers; CNV copy number of the gene is shown as inferred by the HMM applied to normalised coverage data. Amino acid variation is displayed below the dendrogram - black indicates homozygosity and grey heterozygosity. B) Protein model of the N224I mutant *Cyp9k1* docked with deltamethrin, visualised with PyMol. N224I is coloured in green, and can be seen located at the opening of the substrate binding channel of *Cyp9k1*. Inside the channel, the docked Deltamethrin molecule can be observed. C) Diplotype clustering at the *Gste2* locus. D) Protein model of the F120L mutant *Gste2* docked with DDT, visualised with PyMol. F120L is coloured in green, and can be seen located in proximity to the DDT binding domain of *Gste2*. Inside the channel, the docked DDT molecule can be observed.

To explore the potential causative mechanism of these P450 variants, we downloaded the *Cyp9k1* and *Cyp6aa1* protein model predictions from the AlphaFold database^41,42^. After the addition of the heme cofactor, visualisation in PyMol showed that the N224I residue forms part of the opening of the *Cyp9k1* substrate channel (Figure 4B), a small channel which allows organic molecules to pass through to the active site of the enzyme. As Isoleucine is significantly more hydrophobic than Asparagine^43^, this mutation could favour more hydrophobic substrates such as pyrethroid insecticides^44^, which *Cyp9k1* is known to metabolise. In contrast, *Cyp6p3*-D155N is predicted to occur on the surface of the *Cyp6p3* protein somewhat distal from both the active site and substrate channel, and so it may cause indirect effects on metabolism by affecting protein stability or redox partner binding.

At the organophosphate-resistance locus, *Coeaexf*, which contains two alpha-carboxylesterases we find the *Coeaexf-*Dup1 CNV which was previously associated with resistance to pirimiphos-methyl^34^ in *An. gambiae* at 8% frequency. We observe only small clusters of *An. coluzzii* which exhibit low heterozygosity and inter-sample genetic distance, suggesting that any mutations under selection are yet to spread in this species (Supplementary Figure 11).

*Gste* genes are primarily associated with resistance to the organochlorine DDT, however, in previous research they have also been associated with resistance to pyrethroids and organophosphates^45–47^. At the *Gste2* locus, we observed three major selective sweeps (Figure 4C). In line with previous literature, one of these sweeps looks to be driven by the *Gste2*-I114T mutation, and another by the *Gste2*-L119V mutation. Both have previously been associated with resistance to insecticides. Interestingly, the third sweep contains a novel mutation in *An. coluzzii*, *Gste2*-F120L, caused by a G to C substitution (3R:28,598,057). This amino acid variant has previously been reported in *An. maculopennis^48^*, and was computationally modelled in two earlier studies investigating *Gste2*-I114T^46,49^, though it was not followed up on for functional validation. Given its occurrence in other species and its physical proximity to the I114T and L119V residues, this mutation is a strong candidate to be a causal mutation for insecticide resistance. By visualising the *Gste2* enzyme with the F120L mutation (Figure 4D), we confirm that F120L is indeed proximal to the GST active site, as reported in earlier modelling studies^46,50^. To explore the geographical range of *Gste2-*F120L beyond Obuasi, we integrated further data from the Ag1000g phase 3^6^ and a previous GWAS study from West Africa^9^, including whole-genome data from twenty-two sub-Saharan countries (Supplementary Figure 12). These data show that two distinct *Gste2-*F120L mutations exist in sub-Saharan Africa, the aforementioned G>C mutant, as well as a second F120L mutation caused by a G>T nucleotide substitution. The G>C *Gste2-*F120L mutation is found throughout West Africa - in Ghana, Côte d’Ivoire, Mali, Guinea, Burkina Faso and Benin - but only in *An. coluzzii*. In contrast, the G>T F120L mutant is found throughout West to Central Africa - in Mali, Burkina Faso, Ghana, Côte d’Ivoire, Togo, Cameroon and the Central African Republic, but only in *An. gambiae*.

Artisanal gold-mining is associated with the use of mercury and arsenic^51^, toxic elements which can contaminate groundwater and add further selection pressures on malaria vectors in addition to that from insecticides. Based on a univariate generalized linear mixed modeling analysis, we found significant associations between the presence of artisanal gold mining sites and several insecticide resistance alleles after controlling for multiple testing with Benjamini-Hochberg correction (Supplementary Table 4). Among target-site resistance mutations, the *Vgsc* mutation R254K was significantly more common in mining areas (β = 1.12, p = 0.00035), as was the P1874S mutation (β = 0.96, p = 0.00001), whilst we observed that the *Rdl*-A296G mutation was significantly less prevalent (β = -1.14, p < 0.00001). The *Cyp6p3*-E205D mutation was significantly more common (β = 1.38, p < 0.00001). For copy number variants, we found contrasting patterns at the Cyp6aa/p locus, with Cyp6aap_Dup7 more common (β = 1.05, p = 0.00077), and Cyp6aap_Dup10 substantially less common (β = -1.95, p < 0.00001). These findings suggest that artisanal gold mining activities could potentially contribute selection pressures that influence the distribution of insecticide resistance alleles.

## Discussion

In this study, we evaluated the utility of a spatially-explicit sampling framework for enhancing population genomic analyses in malaria vectors. Our approach revealed fine-scale isolation-by-distance in *An. coluzzii* at a high spatial resolution, demonstrating that geographic distance—rather than major ecological factors—primarily drives genetic differentiation at this scale. Through strategic sampling generating a gradient of inter-site distances, we detected significant isolation-by-distance. We found no evidence for genetic structure between ecological strata in our study area. Although ecological stratification could be important for certain aspects of vector surveillance, its impact may emerge only at broader geographic scales or in more ecologically heterogeneous landscapes. Exactly how ecologically-informed sampling frameworks translate into enhanced genomic surveillance is still unclear. Our findings highlight the need for expanded ecological genomics across sub-Saharan Africa to determine the spatial scales at which ecological factors begin to influence population structure and insecticide resistance in malaria vectors.

Artisanal and small-scale gold mining (ASM) can create stagnant pools of water - potential breeding sites for mosquitoes - potentially exacerbating the transmission of malaria^52^. Additionally, heavy metals used in the gold amalgamation process are toxic, and could select for metabolic detoxification processes in malaria vectors that also contribute to insecticide resistance. We did not find evidence for population structure between *An. coluzzii* sampled from ASM villages and non-ASM villages, but the frequency of some resistance variants did differ, perhaps most notably a higher frequency of the important metabolic variant *Cyp6p3-*E205D in ASM village ^40^. The geographical segregation of ASMs to the west of the sampling area in our dataset may have contributed to this effect, however, and so further research is necessary to determine if ASMs are selecting for resistance in vector populations.

The dense sampling enabled the detection of kin in our data. The detection of close genetic relatives, including a pair of full siblings captured 111 meters apart and 3rd-degree relatives in villages 7 km apart, provides evidence of mosquito movement patterns and opens up possibilities for close-kin mark-recapture studies (CKMR). CKMR studies can leverage genetic relationships between individuals to estimate population size and dispersal without the need for physical marking and recapture of mosquitoes. This approach has been successfully applied in other vectors ^53,54^, but remains underutilized in *Anopheles* research, where traditional mark-recapture studies are logistically challenging and yield limited data on natural movement patterns. However, we highlight the complexity of inferring kinship in *An. gambiae*. Large, ancient chromosomal inversions can lead to misleading estimates of relatedness when not accounted for. This phenomenon may affect many published studies, particularly those that employ genome-wide high-density sequencing on non-model organisms. As far as we can see, this has not previously been reported in the literature. In addition, the low number of chromosomes and recombination events in *An. gambiae* compared to other organisms such as humans means that relatives can, by chance, share substantially more or less genetic material than the expectation, complicating traditional kinship inference methods. This chromosomal architecture means that while some siblings may share very large proportions of their genome (appearing more related than typical siblings), others may share much less, potentially being misclassified as more distant relatives. This suggests that future studies of mosquito dispersal may benefit from approaches to kinship estimation that incorporate uncertainty rather than discrete categorisation, particularly when applying close-kin mark-recapture methods.

Our genomic analysis of insecticide resistance revealed both known and novel resistance mechanisms. The continued evolution of the *Vgsc* is evident in the distinct patterns of secondary mutations accompanying the L995F mutation in *An. gambiae* and *An. coluzzii*. The discovery of two independent mutations causing the same amino acid change (F120L) in *Gste2*, with distinct geographic distributions in *An. gambiae* and *An. coluzzii*, provides an example of convergent evolution, a phenomenon continually observed in the field of insecticide resistance^34,55^. The location of this residue near the enzyme’s active site, combined with its independent emergence in multiple species, strongly suggests its functional importance in insecticide resistance. The identification of novel mutations in cytochrome P450s, particularly *Cyp9k1*-N224I and *Cyp6p3*-D155N, adds to our understanding of metabolic resistance. The structural analysis of these mutations suggests different modes of action: N224I putatively affecting substrate specificity through alterations to the substrate channel, while D155N may influence protein stability or partner interactions. We observed particularly contrasting patterns of CNVs between species. At the Cyp6aa/p cluster, *An. coluzzii* shows multiple distinct CNV alleles under selection, while *An. gambiae* exhibits the *Cyp6p3*-E205D mutation with minimal CNV involvement. Similarly, at *Cyp9k1*, *An. gambiae* shows high frequencies of gene amplifications, while *An. coluzzii* displays a strong selective sweep without obvious causative mutations. These patterns suggest that the two species may be evolving different solutions to similar selective pressures. The diversity of resistance mechanisms we observed again indicates that resistance can evolve repeatedly and through various pathways, although the same loci are often involved. This emphasizes the importance of resistance monitoring and management strategies that account for its polygenic nature.

The success of our sampling framework in enabling fine-scale population genetic analyses suggests that similar approaches could be valuable for vector surveillance, though their broader utility remains to be fully established. While our methodology detected subtle population structure, how this translates to improved resistance monitoring requires further investigation. Future studies might combine such spatially explicit sampling with temporal sampling to track the spread of resistance alleles and understand their seasonal dynamics, which could potentially enhance our ability to monitor emerging resistance mechanisms such as *Cyp9k1*-N224I. However, the cost-benefit ratio of implementing such intensive sampling schemes across ecological contexts needs careful evaluation. Additional studies across multiple vector species and varied landscapes will be necessary to determine whether the insights gained justify the additional sampling complexity in routine surveillance programs.

### Materials & Methods

### Sample collection and sequencing

Samples were collected in Obuasi district with CDC light traps, using an ecologically-informed sampling framework^14^. Collections took place from October to December 2018. Mosquitoes were stored in ethanol prior to Illumina 150bp paired-end whole-genome sequencing. Sequencing was performed to a target coverage of 30X and bioinformatic analysis was performed as described previously using a GATK-based workflow following Ag1000g phase 3 protocols^6,56^. In brief paired-end multiplex libraries were processed using Illumina’s DNA preparation protocol and fragmented via Covaris Adaptive Focused Acoustics. Samples were sequenced to ∼30× coverage on Illumina HiSeq 2000 or X platforms yielding an insert size between 100 and 200 bp. Reads were aligned to the AgamP4 reference genome using BWA v0.7.15^57^, with indel realignment and SNP calling performed using GATK v3.7.0^58^. Quality control excluded samples with insufficient coverage (<10× median or <80% genome at 1×) or contamination (≥4.5%), while site filters were developed using colony crosses to identify reliable SNP calls. Genotypes at biallelic SNPs passing filters were phased into haplotypes using a combination of read-backed phasing with WhatsHap v1.0^59^ and statistical phasing with SHAPEIT v4.2.1^60^. The location of informal gold-mining sites were obtained as personal communication from Anglo-Gold Ashanti.

### Genetic Diversity and Landscape Genomics

All population genomic analyses were performed in python with scikit-allel version 1.2.1 and malariagen_data version 10.0.1 unless explicitly stated otherwise. PCA was performed on chromosome 3L, using markers between 15 MB and 44 Mb, to avoid regions of low recombination and known chromosomal inversions. NgsRelate^23^ was used to calculate kinship on biallelic genotypes from across the whole-genome. Linear regression was used to test for isolation-by-distance through F_st_ and the number of shared doubletons with statsmodels^61^. Mantel tests and partial mantel tests were performed in python with code modified from the ecopy package. To test for associations in insecticide resistance alleles with artisanal gold-mining, we performed univariate generalised linear mixed modelling with the python package gpboost using village as a random effect (formula: genotype ∼ mining_status + (1 | village) and controlled for multiple testing with Benjamini/Hochberg error correction with statsmodels fdrcorrection.

### Kinship Simulations

To explore and compare genetic relatedness in human and Anopheline siblings, we performed pedigree simulations with ped-sim v1.4^26^ and analysed the IBD segments and kinship with custom Python scripts. For human simulations, we used a previously published genetic map^62^, for *An. gambiae*, we used previously published maps^4^. We simulated a complex four-generational pedigree structure. Recombination breakpoints were modelled as a poisson process. Each simulation consisted of 10 replicate pedigrees spanning 4 generations with the following structure: Generation 1 contained 4 branches with one individual printed per branch. Generation 2 expanded to 4 branches with 2 individuals printed per branch. Generations 3 and 4 each contained 12 branches with 2 individuals printed per branch. 800 individuals were generated across all replicates for each species.

### Genome-wide Selection Scans

We performed genome-wide selection scans with the H12 statistic^32^. This statistic is a measure of haplotype homozygosity, and captures is particularly powerful in detecting soft selective sweeps, or where there are multiple distinct haplotypes under selection at the same locus. We then took each cohort and used malariagen_data to compute H12 in 1500 SNP stepping windows. To investigate haplotype sharing between cohorts, we used the H1X statistic, which measures haplotype sharing between two cohorts^33^.

### Diplotype Clustering

We extracted diplotypes from windows around each putative resistance locus and clustered the diplotypes with complete-linkage hierarchical clustering. Diplotypes, sometimes referred to as multi-locus genotypes, are stretches of diploid genotypes. For each site, we then find the pairwise difference between the allele counts for each individual in the pair. We then use city-block (Manhattan) distance as the distance metric and complete linkage. We also determined which non-synonymous variants were present on each diplotype and plotted this data alongside sample heterozygosity over the region, and the number of extra copies of each gene in the region.

### Sample Heterozygosity

We included heterozygosity in the diplotype clustering plots, to permit easy identification of diplotype clusters homozygous for a selective sweep, as opposed to diplotype clusters which contain two separate haplotypes under selection. For a given diplotype region, we calculated the observed heterozygosity for each variant and then averaged these per-variant heterozygosity values to get a single value of heterozygosity for the region for each individual sample.

### CNV Detection

To detect CNV alleles at the insecticide resistance loci investigated in this study, we used the Ag1000g coverage-based CNV calls, which applies a hidden Markov model (HMM) to normalized sequencing coverage to estimate the copy number state in 300 bp genomic windows as previously described^5^. In brief; sequencing coverage is first normalized to account for variation in local nucleotide composition. A HMM is then applied to the normalized, filtered, coverage data. The HMM contains 13 hidden states, representing copy numbers from 0 to 12 in increments of 1, allowing the detection of up to a 6-fold amplification of a genetic region, or 10 extra copies.

### Protein Modelling

We downloaded predicted protein models from the AlphaFold database^41,42^, selecting the longest transcript present, to avoid predictions of truncated transcripts. For P450s, we then used ChimeraX^63^ to superimpose the PDB model of Homo Sapiens *Cyp3a4* (PDB: 1TQN), in order to transplant the Heme cofactor into our predicted model. We visualised the protein models with PyMol^64^.

## Code and Data availability

Code for this manuscript is stored here https://github.com/sanjaynagi/obuasi_ms. A snakemake workflow to infer kin with NgsRelate is here https://github.com/sanjaynagi/ag3-relatedness. Read data for the samples analysed in this study is stored in the ENA. Accession codes are stored in Supplementary Table 1.

## Acknowledgments

Research reported in this publication was supported by the Medical Research Council (MR/P02520X/1 and MR/T001070/1). MR/P02520X/1 is part of the EDCTP2 programme supported by the European Union. The work herein was also supported by the National Institute of Allergy and Infectious Diseases of the National Institutes of Health under Award Number R01AI116811. The content is solely the responsibility of the authors and does not necessarily represent the official views of the National Institutes of Health. MJD is supported by a Royal Society Wolfson Fellowship (RSWF\FT\180003).

