## Supplemental Text and Figures for "Spatially-explicit genomics of *An. gambiae s.l* uncovers fine-scale population structure and mechanisms of insecticide resistance"

### Obuasi supplement

#### Supplementary Text 1

Suppl. Figure 2A shows *An. coluzzii*, in which two pairs of outliers appear distinct from the other individuals. Interestingly, each pair of outliers were sampled from the same village. Removal of these four outliers (Suppl. Figure 2B) results in four new samples being pulled out by the PCA as outliers, which previously clustered with the rest of the *An. coluzzii* individuals. In *An. gambiae*, we observed a similar pattern (Supplementary Figure 2), with two outlying samples. However, the removal of these two samples resulted in a PCA which was much more clearly without structure (Suppl. Figure 2D). Overall, the principal components analysis suggests that there is little variation within *An. coluzzii* or *An. gambiae s.s*, in agreement with earlier findings that this region of West Africa is relatively homogeneous (Ag1000G, 2020).

#### Supplementary Text 2

Chromosomal inversions in *Anopheles gambiae* are known to be ancient - for example, the 2La inversion predates speciation into the gambiae complex, and therefore genetic differentiation between opposing karyotypes is very large - greater than the differentiation found between species (ref). To evaluate how chromosomal inversions affect the inference of relatedness, chromosomal inversions were scored with a modified version of compkaryo (Love *et al.*, 2019). Supplementary table 2 shows karyotype frequencies for each sample site in Obuasi for the 2La and 2Rb inversions (*An. coluzzii*). Supplementary Figure 3 shows the results of the kinship analysis with inclusion of chromosomal inversion regions. Three clusters of KING-robust values can be observed which, relate to the pairwise 2La karyotype concordance. The negative values centering around -0.1 correspond to comparisons in which both individuals were homozygous for alternative 2La inversion karyotypes, the cluster centred around 0 contains primarily pairwise comparisons between homozygotes and heterozygotes, and the cluster at ≈0.075 contains pairwise comparisons with identical 2La karyotypes (Supplementary Figure 3). Based on the thresholds set by (Manichaikul *et al.*, 2010), this would result in almost all intra-karyotypic comparisons being defined as 3rd-degree relatives. This suggests that the KING estimator, and any estimator based on levels of linkage disequilibrium may have poor performance at lower degrees of relatedness when inversions are present in the data. We therefore recalculated kinship while masking the regions of the 2La and 2Rb. Supplementary Figure 4 shows the results from this analysis, with the data exhibiting a single cluster of KING values centred around 0.

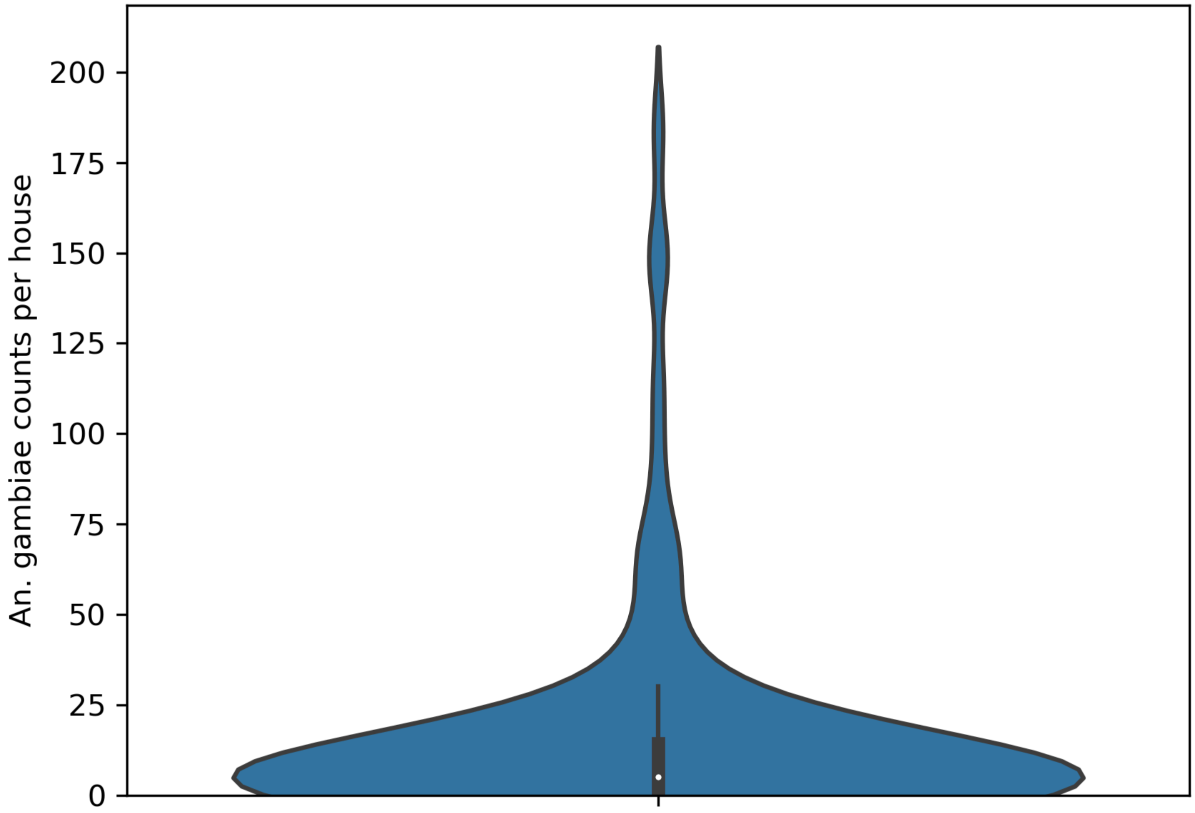


**Supplementary Figure 1. A violin plot of counts per house of *An. gambiae* during the sample collections.** Five sample collections in each house were performed over a 9 week period.

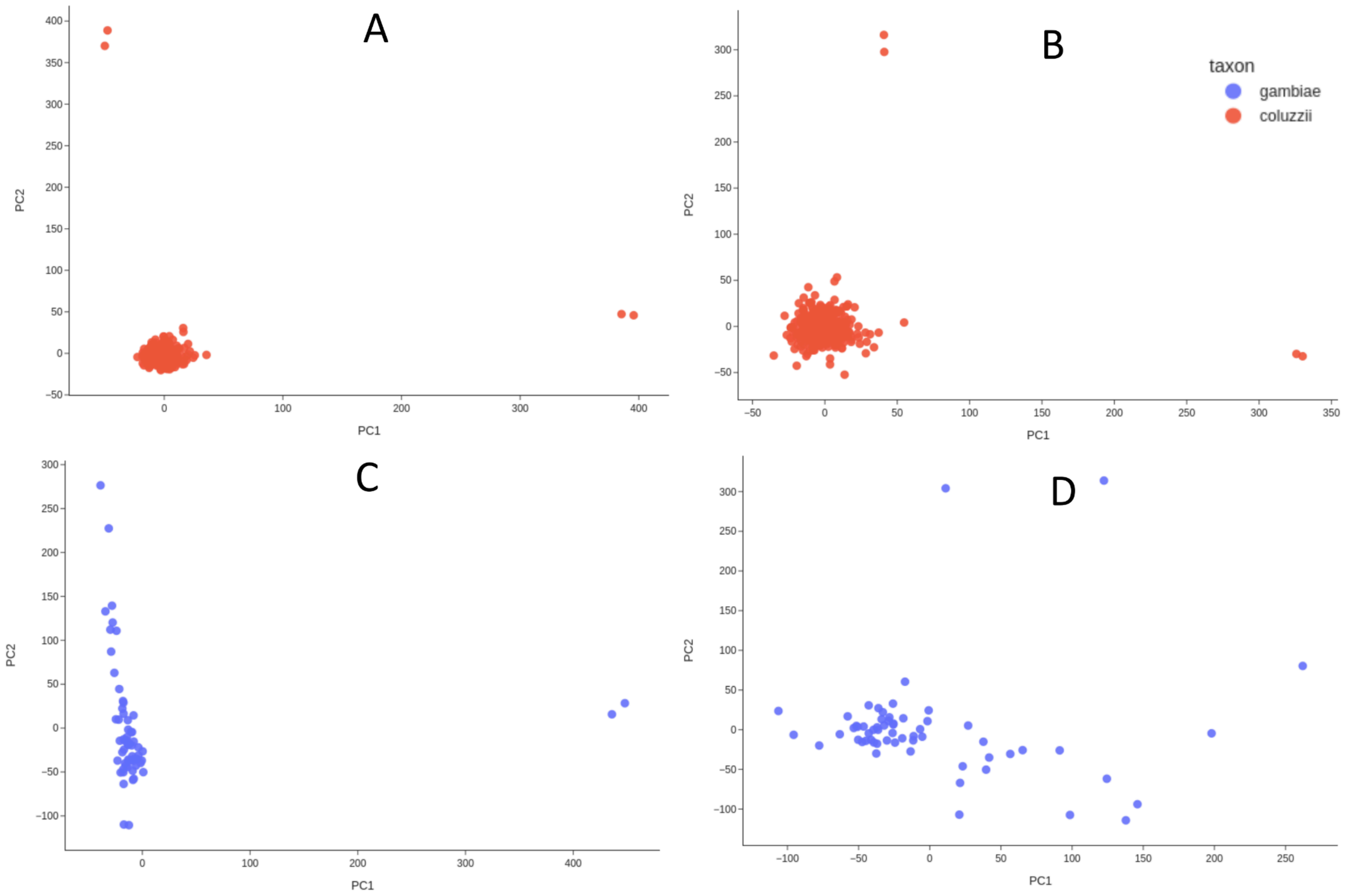


**Supplementary Figure 2. Principal components analysis on chromosomal arm 3L, 15 Mb - 44 Mb.** A) *An. coluzzii*, B) *An. coluzzii* after removal of outliers C) *An. gambiae* D) *An. gambiae* after removal of outliers.

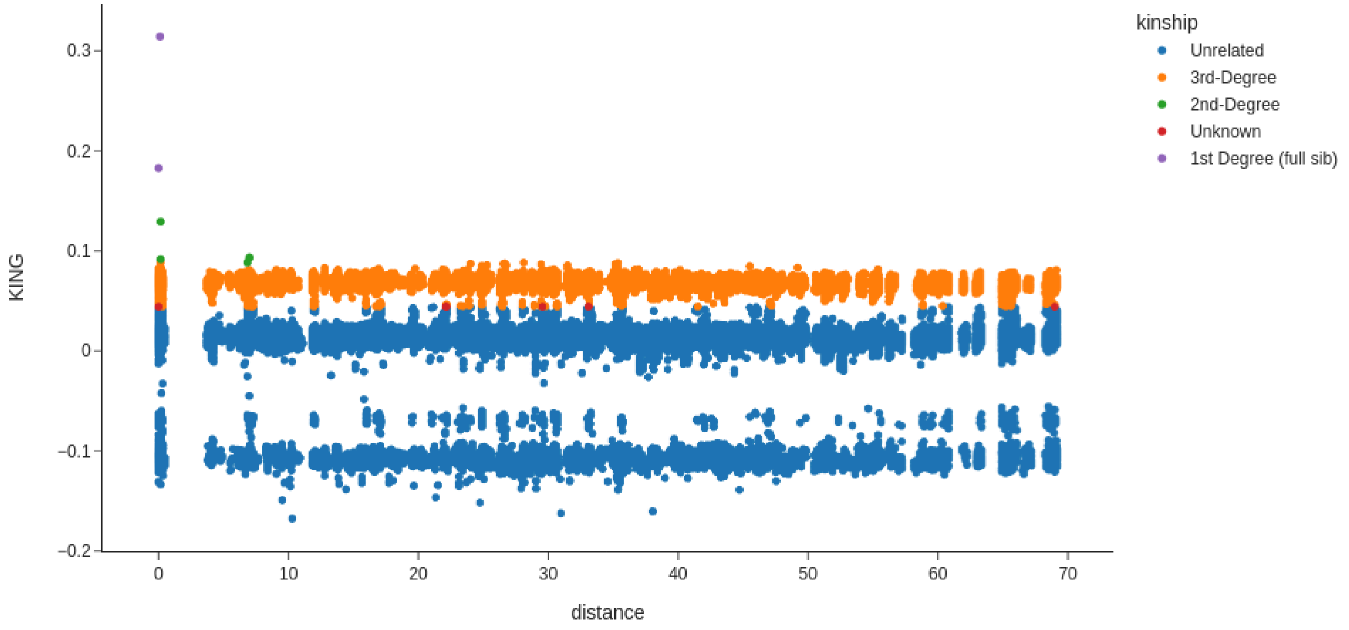


**Supplementary Figure 3. Pairwise relatedness (KING-robust) between samples plotted against geographic distance in kilometres, without masking for chromosomal inversions.** Points are coloured depending on the thresholds advised by the KING authors. Pairwise relatedness is calculated using NgsRelate on biallelic SNPs across the whole genome. 1st Degree kin equate to full sibling or parent-child relationships, 2nd Degree is aunt, uncle, grandparent, grandchild, niece, nephew, or half-sibling relationship.

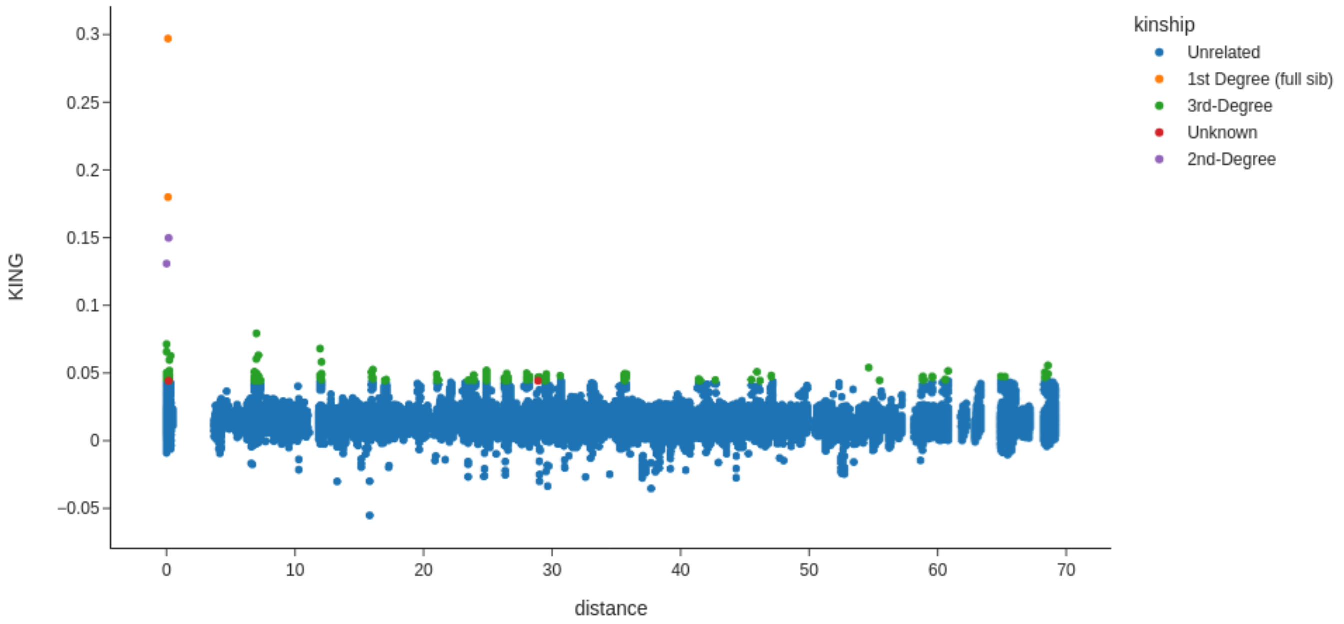


**Supplementary Figure 4. Pairwise relatedness (KING-robust) between samples plotted against geographic distance in kilometres, while masking for chromosomal inversions.** Points are coloured depending on the thresholds advised by the KING authors. Pairwise relatedness is calculated using NgsRelate on biallelic SNPs across the whole genome. 1st Degree kin equates to full sibling or parent-child relationships, 2nd Degree is aunt, uncle, grandparent, grandchild, niece, nephew, or half-sibling relationship.

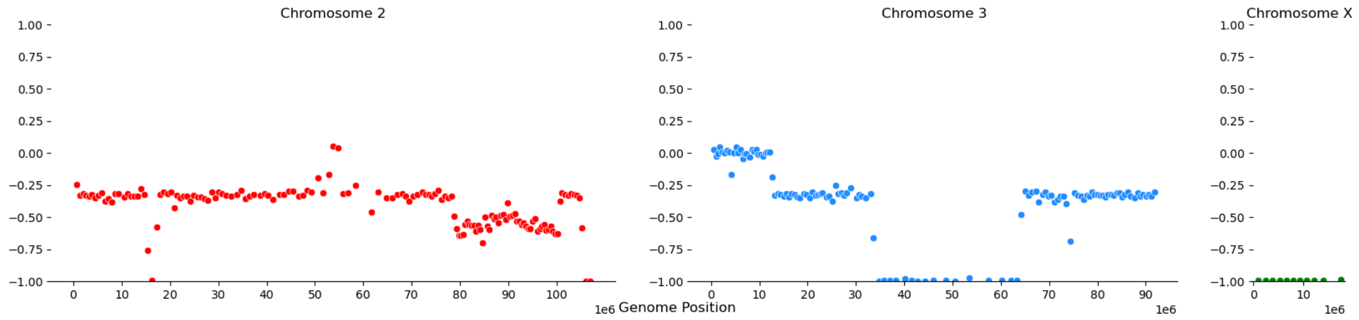


**Supplementary Figure 5.** Plots of Hudson's $F_{st}$ across the genome of two individuals designated as full siblings as per the KING statistic (WA-2361, WA-2363, Female *An. coluzzii* collected in Odumto, in separate houses 111m apart). $F_{st}$ was calculated in 10,000 bp stepping windows.

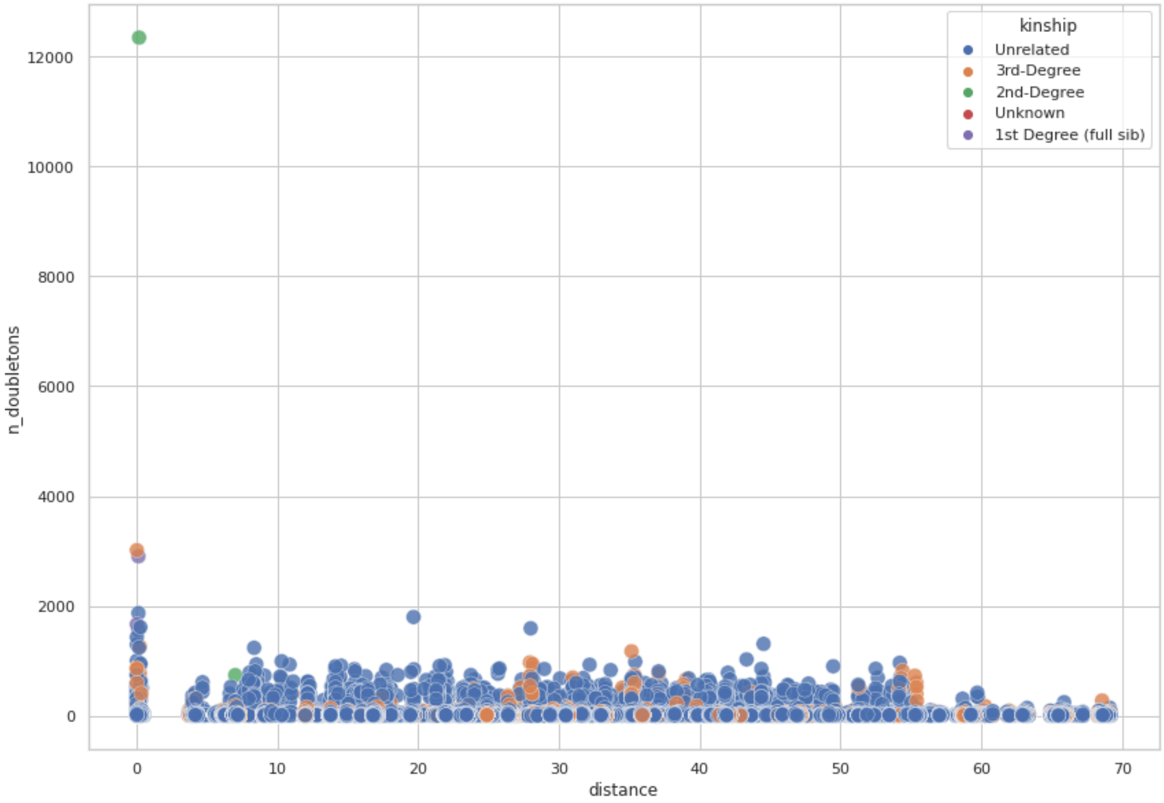


**Supplementary Figure 6.** The number of doubletons vs geographic distance between individual samples, coloured by kinship category.

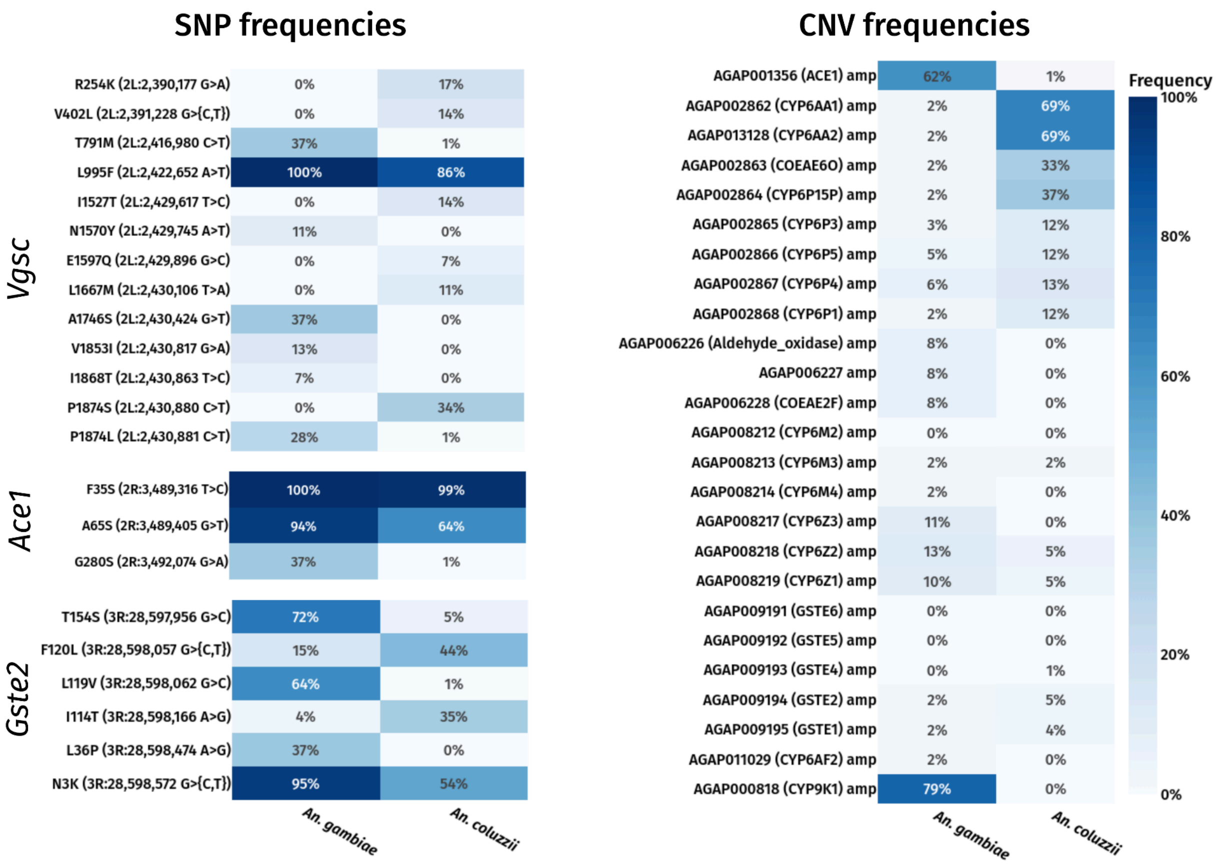


**Supplementary Figure 7. Allele frequencies of SNP and CNV variants.** We calculated the frequencies of a priori SNPs and CNVs involved in insecticide resistance in both *An. gambiae* and *An. coluzzii*.

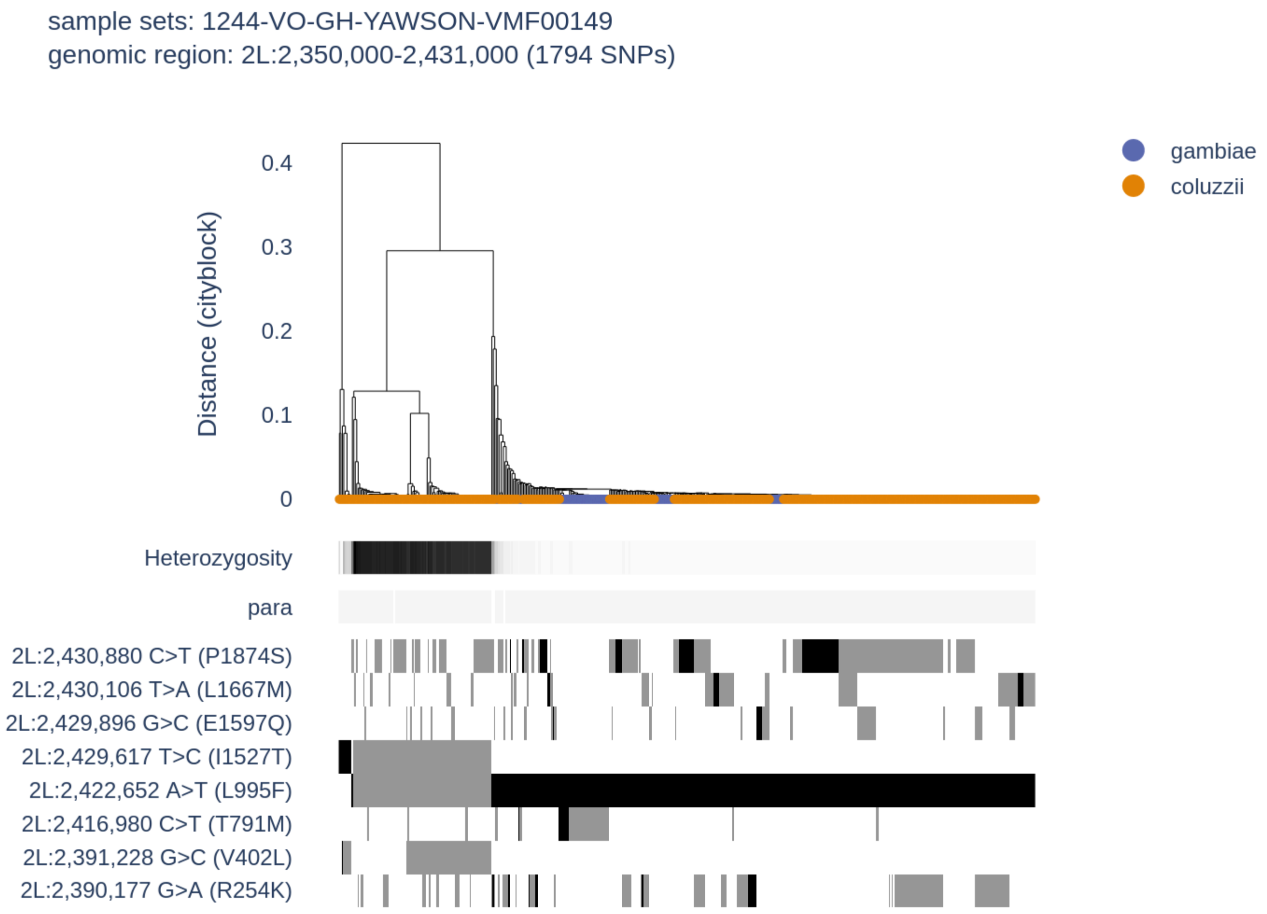


**Supplementary Figure 8.** Diplotype clustering over the *Vgsc* region. We calculate pairwise distance between diplotypes. Each column in the figure is a diplotype ordered by the dendrogram by hierarchical clustering, using genetic distance based on city-block (Manhattan) distance and complete linkage. The leaves of the dendrogram are coloured by the species of the individual to which they belong. Note that due to overlapping points, not all dendrogram leaves can be seen. Underneath the dendrogram, the heterozygosity and CNV copy number of an individual are displayed as horizontal bars. Heterozygosity; individual-level heterozygosity was calculated as an average over all SNPs in the locus. Clusters with low sample heterozygosity or inter-sample genetic distances of zero are indicative of a selective sweep. CNV copy numbers; CNV copy number of the gene is shown as inferred by the HMM applied to normalised coverage data. Amino acid variation is displayed below the dendrogram - black indicates homozygosity and grey heterozygosity.

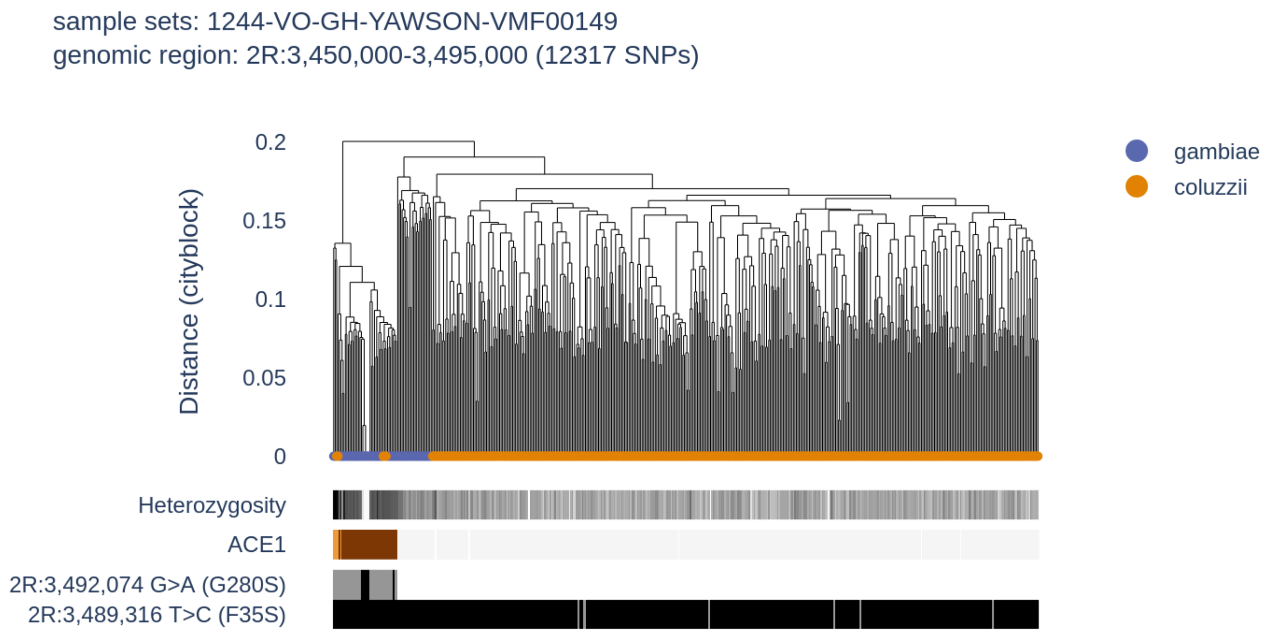


**Supplementary Figure 9.** Diplotype clustering over the *Ace1* region. We calculate pairwise distance between diplotypes. Each column in the figure is a diplotype ordered by the dendrogram by hierarchical clustering, using genetic distance based on city-block (Manhattan) distance and complete linkage. The leaves of the dendrogram are coloured by the species of the individual to which they belong. Note that due to overlapping points, not all dendrogram leaves can be seen. Underneath the dendrogram, the heterozygosity and CNV copy number of an individual are displayed as horizontal bars. Heterozygosity; individual-level heterozygosity was calculated as an average over all SNPs in the locus. Clusters with low sample heterozygosity or inter-sample genetic distances of zero are indicative of a selective sweep. CNV copy numbers; CNV copy number of the gene is shown as inferred by the HMM applied to normalised coverage data. Amino acid variation is displayed below the dendrogram - black indicates homozygosity and grey heterozygosity.

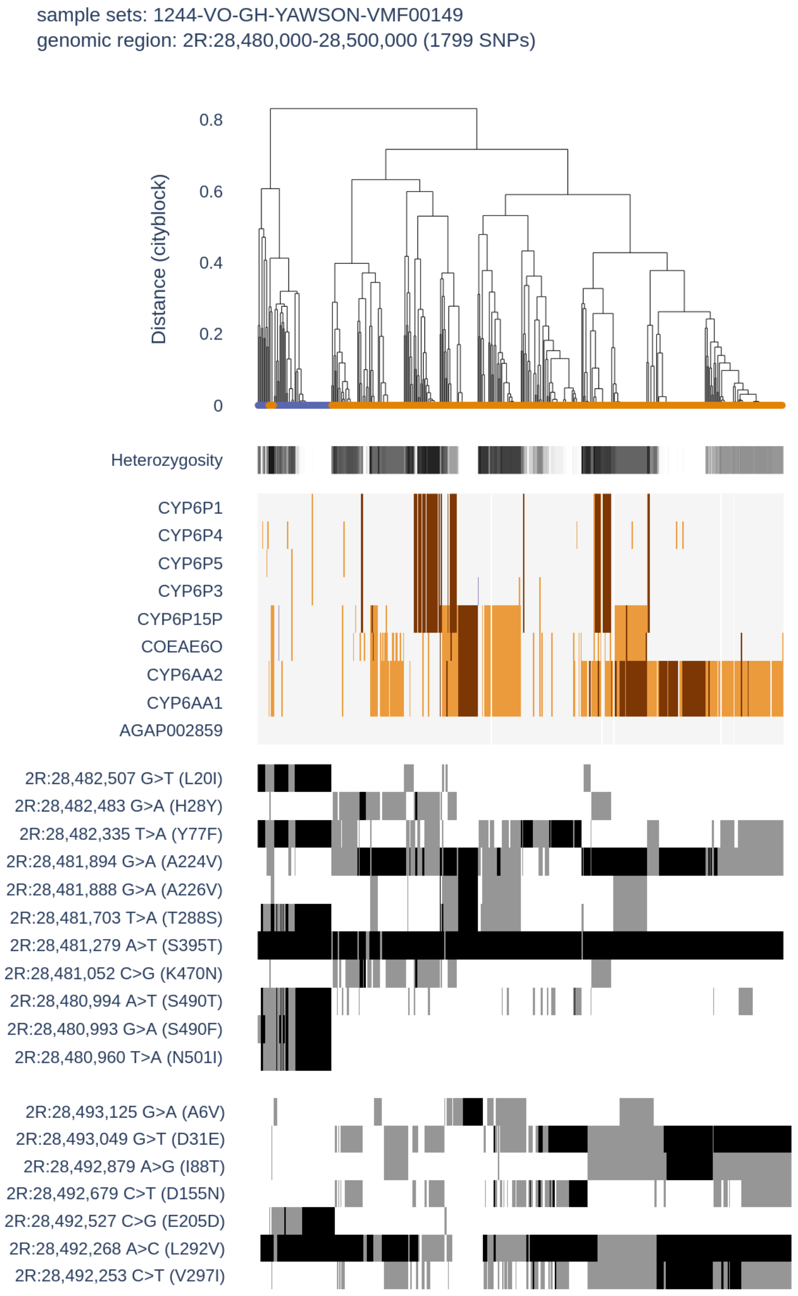


**Supplementary Figure 10.** Diplotype clustering over the *Cyp6aa1* and *Cyp6p3* region. We calculate pairwise distance between diplotypes. Each column in the figure is a diplotype ordered by the dendrogram by hierarchical clustering, using genetic distance based on city-block (Manhattan) distance and complete linkage. The leaves of the dendrogram are coloured by the species of the individual to which they belong. Note that due to overlapping points, not all dendrogram leaves can be seen. Underneath the dendrogram, the heterozygosity and CNV copy number of an individual are displayed as horizontal bars. Heterozygosity; individual-level heterozygosity was calculated as an average over all SNPs in the locus. Clusters with low sample heterozygosity or inter-sample genetic distances of zero are indicative of a selective sweep. CNV copy numbers; CNV copy number of the gene is shown as inferred by the HMM applied to normalised coverage data. Amino acid variation is displayed below the dendrogram - black indicates homozygosity and grey heterozygosity.

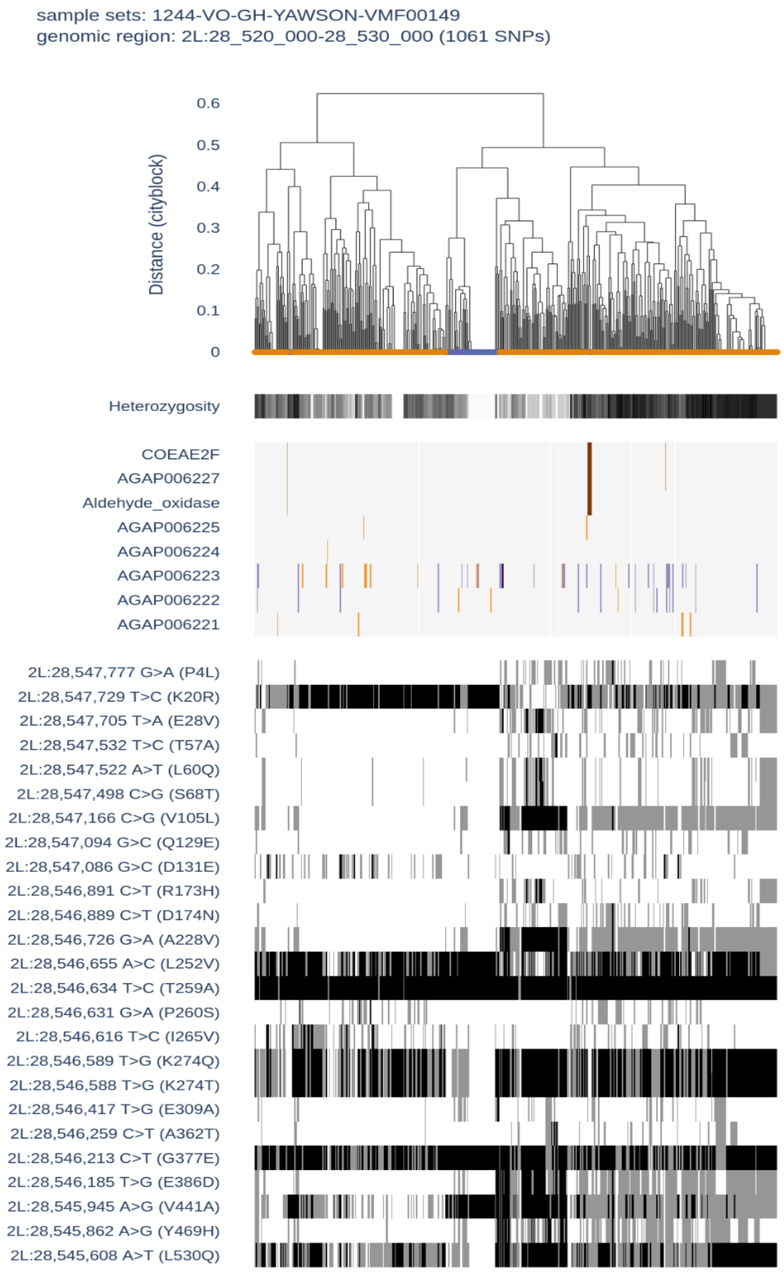


**Supplementary Figure 11.** Diplotype clustering over the *Coeae1f* region. We calculate pairwise distance between diplotypes. Each column in the figure is a diplotype ordered by the dendrogram by hierarchical clustering, using genetic distance based on city-block (Manhattan) distance and complete linkage. The leaves of the dendrogram are coloured by the species of the individual to which they belong. Note that due to overlapping points, not all dendrogram leaves can be seen. Underneath the dendrogram, the heterozygosity and CNV copy number of an individual are displayed as horizontal bars. Heterozygosity; individual-level heterozygosity was calculated as an average over all SNPs in the locus. Clusters with low sample heterozygosity or inter-sample genetic distances of zero are indicative of a selective sweep. CNV copy numbers; CNV copy number of the gene is shown as inferred by the HMM applied to normalised coverage data. Amino acid variation is displayed below the dendrogram - black indicates homozygosity and grey heterozygosity.

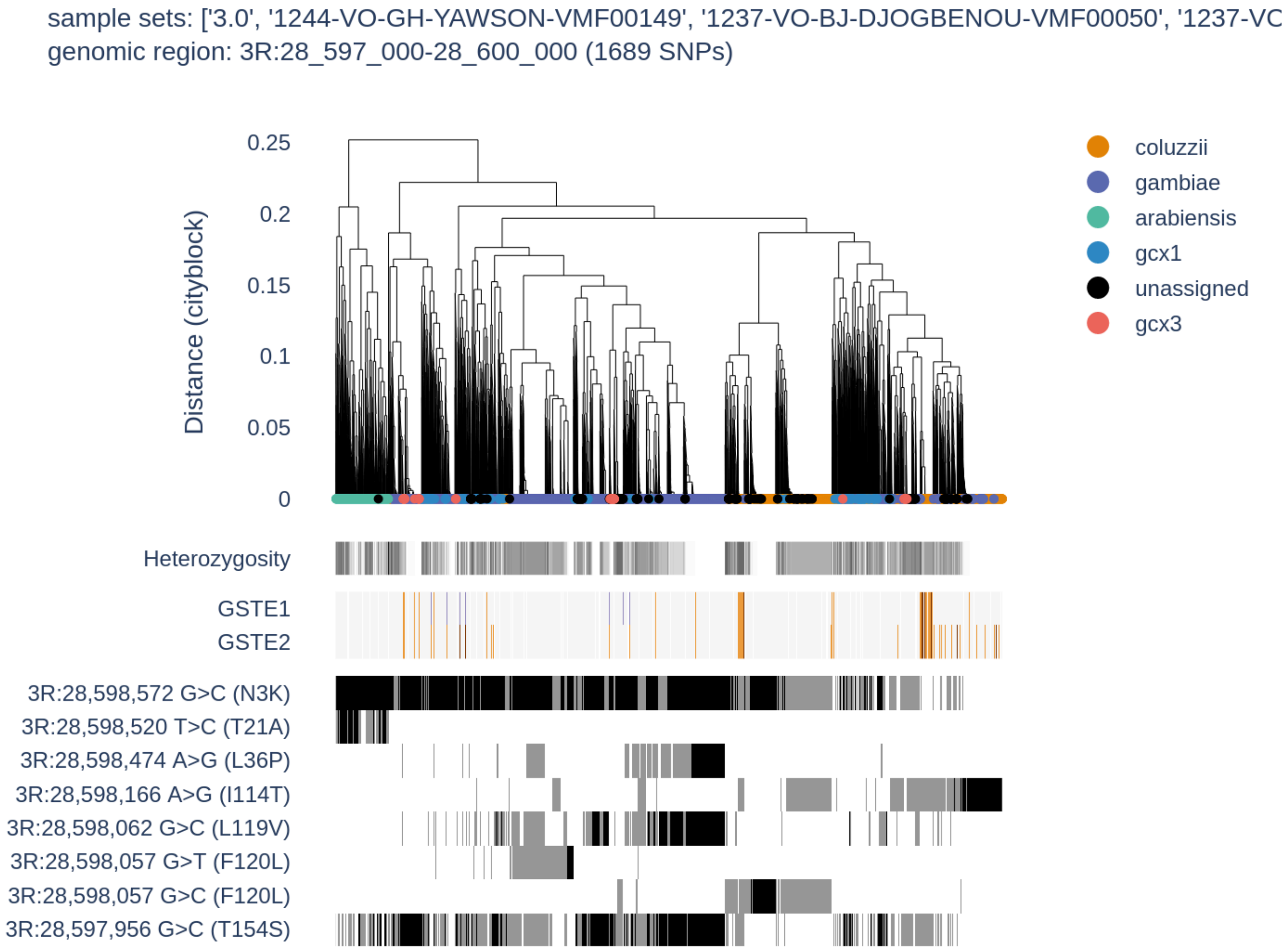


**Supplementary Figure 12. A)** Diplotype clustering over the *Gste2* region in Obuasi data **B)** In data from Obuasi, Ag1000g Phase 3, and a recent GWAS study in West Africa. We calculate pairwise distance between diplotypes. Each column in the figure is a diplotype ordered by the dendrogram by hierarchical clustering, using genetic distance based on city-block (Manhattan) distance and complete linkage. The leaves of the dendrogram are coloured by the species of the individual to which they belong. Note that due to overlapping points, not all dendrogram leaves can be seen. Underneath the dendrogram, the heterozygosity and CNV copy number of an individual are displayed as horizontal bars. Heterozygosity; individual-level heterozygosity was calculated as an average over all SNPs in the locus. Clusters with low sample heterozygosity or inter-sample genetic distances of zero are indicative of a selective sweep. CNV copy numbers; CNV copy number of the gene is shown as inferred by the HMM applied to normalised coverage data. Amino acid variation is displayed below the dendrogram - black indicates homozygosity and grey heterozygosity.
